# Borrowing data from other populations to forecast epidemic size

**DOI:** 10.64898/2026.09.18.752593

**Authors:** Sam Paplauskas

## Abstract

Forecasting infectious disease dynamics is important for ecological management and emerging disease preparedness, yet many systems lack the long-term datasets required to develop reliable predictions. Spatial replication may provide an alternative by allowing information to be shared among populations, but the extent to which this improves forecasting remains unclear. Here, I used epidemic and temperature data from 20 replicated semi-natural *Daphnia*–parasite pond populations monitored across four seasons (80 epidemics) to test whether information from multiple populations improves forecasts and whether environmental similarity identifies informative populations. I forecasted disease prevalence, infected host density, and healthy host density using a benchmark model, autoregressive integrated moving average (ARIMA), and time-series regression models. Models were trained using focal-population data alone, mean dynamics across other populations, or temperature-weighted mean dynamics from environmentally similar populations. Epidemics showed strong seasonal structure, with prevalence and infected host density generally peaking during the warmest period of the season. However, no forecasting approach consistently outperformed others across all variables. Temperature improved forecasts of prevalence and infected host density but not healthy host density. Multi-population training improved forecasts in some cases, particularly for ARIMA models, whereas temperature-weighted averaging provided no additional benefit over simple averaging. Regression models consistently produced the most accurate forecasts of healthy host density. These results show that borrowing information among populations can improve forecasts under specific conditions but is not universally beneficial, highlighting the importance of matching forecasting approaches and data sources to the ecological processes being predicted.

## Introduction

Infectious disease epidemics can have profound consequences for the ecology, evolution and conservation of natural populations by reducing host survival, reproduction and population size, while also influencing community structure and ecosystem processes (Altizer et al., 2006; Tompkins et al., 2011). Accurately quantifying and predicting epidemic size is therefore a central goal of disease ecology because it enables researchers and managers to anticipate disease impacts and implement timely interventions. Epidemic size is commonly measured as disease prevalence, the proportion of hosts infected within a population (Jennelle et al., 2007). However, prevalence is not always the most informative metric. The density of infected hosts (the number of infected individuals per unit area or volume) may provide a more direct measure of transmission risk, but transmission also depends on the local availability of susceptible hosts. Thus, the density of healthy hosts may provide important information about the pool of hosts available for infection, as well as being relevant to conservation and population management because it reflects the number of susceptible or surviving individuals. Together, these complementary measures provide a more complete description of epidemic dynamics and the ecological consequences of disease.

Forecasting remains relatively underdeveloped in wildlife epidemiology compared with human epidemiology, where disease forecasting systems are more routinely used for public health decision-making (Lutz et al., 2019). Where forecasting has been applied in wildlife disease systems, approaches have generally relied on historical observations from the same population, using time-series methods to identify temporal patterns in epidemic dynamics and, in some cases, incorporating environmental drivers such as temperature (Gao et al., 2012; Hii et al., 2012; Hu et al., 2006). However, long-term disease time series are difficult and costly to collect in wildlife populations, and are particularly unlikely to be available for emerging diseases or newly established host–parasite associations. This limitation highlights the need for alternative approaches that can generate forecasts when historical data from a focal population are limited or absent.

Here, I propose that spatial variation among populations can provide an alternative source of predictive information, allowing data from other populations to supplement or substitute for missing temporal information from a focal population. If populations exposed to similar environmental conditions exhibit similar epidemic dynamics, then environmentally similar populations may provide informative analogues for forecasting future epidemics. Temperature is particularly promising in this context because it is a key driver of many host–parasite interactions and can often be measured across populations even when disease observations are unavailable (Alonso et al., 2011; Auld & Brand, 2017b; Beckley et al., 2016; Bravo et al., 2020; Groner et al., 2018, 2021; Krauer et al., 2021; Ruiz-Moreno et al., 2012; Schaaf et al., 2017; Susi et al., 2017; Swinford & Anderson, 2021; Thoirain et al., 2007).

Existing time-series forecasting approaches provide a framework for incorporating environmental information into predictions by modelling temporal patterns in disease dynamics and their relationships with environmental drivers. Two commonly used approaches include autoregressive integrated moving average (ARIMA) and time-series regression models. ARIMA models make inferences based on underlying patterns of temporal autocorrelation and, although they have previously been used in disease forecasting without environmental predictors (Allard, 1998; Helfenstein, 1991), they can be extended to incorporate additional information, such as seasonal patterns or environmental covariates, into epidemic predictions (Hyndman & Athanasopoulos, 2021). Similarly, time-series regression models incorporate predictor variables that explain variation in disease dynamics (Hyndman & Athanasopoulos, 2021) and have frequently been used to forecast vector-borne diseases, such as malaria and dengue, using environmental factors including temperature and rainfall (Gao et al., 2012; Hii et al., 2012; Hu et al., 2006). These approaches provide a potential framework for incorporating spatial information into disease forecasts by using environmental similarity among populations as a predictor of epidemic dynamics. For example, populations with similar temperature regimes may provide informative data for predicting disease dynamics in a focal population, particularly when historical time-series data from that population are limited.

Here, I used epidemic and temperature data collected from 20 replicated semi-natural *Daphnia*-parasite pond populations over four seasons (80 epidemics) to test whether information from multiple populations can improve forecasts of epidemic dynamics and whether environmental similarity can be used to identify informative populations for prediction. I predicted three measures of epidemic size (disease prevalence, infected host density and the density of healthy hosts) using three time-series forecasting approaches: a benchmark model, an ARIMA model and a time-series regression model. To evaluate whether spatial information can compensate for limited temporal data, I trained models using three alternative datasets: data from only the focal population (single population models), average epidemic dynamics across other populations (average population models), and temperature-weighted averages of other populations, where populations with more similar temperature dynamics to the focal population contributed more strongly to predictions (weighted average population models). I hypothesised that (i) ARIMA and time-series regression models would outperform benchmark models because they can capture more complex temporal patterns, and (ii) models incorporating information from multiple populations would improve forecasts relative to models trained only on the focal population because they increase the amount of available information. However, my results were more nuanced, with model performance varying according to both the forecasting approach and the type of data used for training.

## Methods

### Summary

The dataset corresponding to four years of seasonal epidemics in 20 *Daphnia*-microparasite semi-natural pond populations was used to test whether borrowing data from other populations can outperform forecasts based on isolated populations. As outlined at the end of the introduction, this involved predicting three variables (disease prevalence, infected host density and the density of healthy hosts) using three types of time-series forecasting models (benchmark, ARIMA and time-series regression) that were trained on three subdivisions of my dataset (single population, average population and population data weighted by temperature similarity).

For each of my three forecast variables, I address the following questions (Q1-3):

Q1: What is the temporal (seasonal) pattern of the forecast variable?
Q2: In general, does forecast error vary by the class of forecast model, the data used to train a model and the temperature of the focal population?
Q3: Is forecast error consistently lower for specific combinations of forecast model and training data?

### Study system

In this study, I focused on the *Daphnia magna–Pasteuria ramosa* host–parasite system. *D. magna* is a small freshwater crustacean that naturally occurs with the obligate sterilizing bacterial microparasite *P. ramosa*. In the wild, *D. magna* populations experience regular annual epidemics of *P. ramosa*, with epidemic dynamics emerging from interactions between host demography, parasite transmission, and environmental conditions (Ebert, 2005; Decaestecker et al., 2007). Hosts become infected when they ingest environmentally persistent parasite spores during filter feeding. Following infection, *P. ramosa* sterilizes hosts and produces large numbers of spores that are released after host death and accumulate in pond sediments, creating a reservoir of infectious stages that drives future transmission (Ebert, 2005; Decaestecker et al., 2004; Decaestecker et al., 2007). Consequently, changes in host population size influence epidemic dynamics by affecting the abundance of infected hosts and the production of environmental parasite stages, even though transmission occurs through exposure to environmental spores rather than direct host contact.

Epidemics typically begin as host densities increase in spring, creating conditions favourable for parasite transmission, before prevalence changes throughout the summer as host population dynamics, parasite accumulation, and environmental conditions interact to shape epidemic trajectories (Ebert, 2005; Decaestecker et al., 2007). Seasonal declines in host density and environmental changes contribute to reductions in transmission during autumn, and parasites may disappear from the active host population during winter, although spores can persist in sediments and contribute to future epidemics (Ebert, 2005; Decaestecker et al., 2007). Temperature may further influence epidemic dynamics by affecting host and parasite traits, including host susceptibility, parasite development, and the timing of infection processes, making it a potentially informative predictor of disease dynamics in this system (Mitchell et al., 2005; Auld & Brand, 2017).

### Pond experiment

The pond experiment was part of a broader experimental design investigating how host genetic diversity and environmental variation influence *Daphnia magna–Pasteuria ramosa* coevolutionary trajectories (Paplauskas et al., 2021). Here, I use the resulting multi-year epidemic time series to evaluate the predictability of disease dynamics under different forecasting approaches. A full description of the experimental design is provided below, with details relevant to the generation of the forecasting dataset retained.

To start with, replicate lines of the 12 genotypes of *Daphnia magna* were maintained in the laboratory in a state of clonal reproduction for three generations to reduce variation due to maternal effects. There were five replicates per genotype; each replicate consisted of five *Daphnia* kept in 200 ml of artificial medium (Klüttgen et al., 1994) modified using 5% of the recommended SeO2 concentration (Ebert et al., 1998). Replicate jars were fed 5.0 ABS of *Chlorella* vulgaris algal cells per day (where ABS is the optical absorbance of 650 nm white light by the *Chlorella* culture). *Daphnia* medium was changed three times per week and three days prior to the start of the pond experiment. On the day that the pond experiment commenced, 1–3 day old offspring were pooled according to host genotype. Ten offspring per genotype were randomly allocated to each of the 20 ponds (giving a total of 120 *Daphnia* per pond). This common starting population composition was established as part of a broader experimental design used to investigate evolutionary trajectories under different environmental conditions (Paplauskas et al., 2021).

Each pond consisted of a 0.65 m tall 1000 litre PVC tank filled with rainwater. The ponds were set to different depths into the ground and experienced different temperature profiles (Auld & Brand, 2017b). In addition, six of the ponds experienced a weekly mixing treatment where mixed ponds were stirred once across the middle and once around the circumference with a 0.35 m^2^ paddle submerged halfway into the pond (the exception to this was on the first day of the experiment, when all ponds experienced the mixing treatment to ensure hosts and parasites were distributed throughout the ponds).

The experiment began on the 2nd April 2015 (Julian day 98), when 120 *Daphnia* (10 *Daphnia* x 12 genotypes) and 1 x 10^8^ *Pasteuria* spores from the mastermix were added to each of the 20 ponds. This inoculum concentration was selected to ensure reliable parasite establishment across replicate ponds under semi-natural conditions, where environmental variation and stochastic processes can otherwise limit infection establishment. Although the initial parasite exposure was higher than would typically be observed in natural populations, all ponds received the same standardized inoculum, allowing subsequent differences in epidemic dynamics to be compared among replicate populations. The mastermix comprised *Pasteuria ramosa* spores propagated using 21 separate *Daphnia* genotypes exposed to sediment from their original pond (Kaimes, Scottish Borders, UK, Auld & Brand, 2017).

Seasonal epidemics were tracked for the next four years between April 2015 and November 2018. This involved weekly estimates of parasite prevalence, diseased adult density, healthy adult density, and pond temperature between either April or May and November each year (Auld & Brand, 2017b; Paplauskas et al., 2021). Host densities were estimated from pond net samples, where individuals collected within a known sampled volume were counted and scaled to the total pond water volume to estimate population-level densities.

### Preparation of time-series for forecasting

Following the pond experiment, I processed the resulting longitudinal data into the format required for forecasting. Each pond was represented by a multivariate time series (MTS), consisting of weekly measurements of environmental and population-level variables collected over four years. Specifically, each MTS included pond temperature, an estimate of the number of healthy adult *Daphnia*, and an estimate the number of diseased adult *Daphnia*. From these original time series, I calculated disease prevalence (the proportion of adult hosts that were infected), healthy adult density, and diseased adult density. Density variables were natural log-transformed prior to model fitting to reduce skew and account for differences in population size among ponds.

Weeks were indexed using the POSIX %W week numbering convention, in which Monday is defined as the first day of the week. Under this convention, week indices range from 00 to 53, where Week 00 comprises the days preceding the first Monday of the calendar year and Week 01 begins on the first Monday. Week 53 occurs only in years in which an additional week is required. These values represent week indices used for temporal aggregation and should not be interpreted as indicating that every year contains 54 calendar weeks.

Where multiple observations were recorded within the same week, values were averaged to generate a single weekly estimate. Sampling occurred seasonally, during the period when ponds were monitored (spring–autumn), and therefore each annual time series contained a defined number of weekly observations corresponding to this monitoring period. The first three years of data had a seasonal frequency of 31 weekly observations, which were used as the training data, while the final year had a seasonal frequency of 26 weekly observations and was used as the forecasting evaluation period. Weeks outside the monitoring period were not considered missing observations because no sampling was conducted during these periods. Within the monitoring period, missing values resulting from occasional missed sampling events were linearly interpolated to maintain a consistent time-series structure. Missing observations requiring interpolation were infrequent and were limited to occasional short gaps within the monitoring period. However, where three or more consecutive weekly observations were missing, values were retained as NA rather than interpolated.

### Q1: Seasonal dynamics of epidemiological variables and temperature

To determine whether the variables selected for forecasting exhibited consistent seasonal patterns (Q1), I first characterised their temporal dynamics across the four years of the experiment. Specifically, I examined whether disease prevalence, infected host density, and healthy host density followed repeatable seasonal trajectories, and whether these patterns corresponded with seasonal variation in temperature. Seasonal trajectories of temperature and epidemiological variables were plotted together and compared by visual inspection. This exploratory analysis provided an assessment of whether temperature and population-level disease metrics exhibited predictable seasonal structure, which is important because temporal forecasting approaches rely on the presence of recurring patterns that can be leveraged to predict future dynamics. In addition, identifying the extent to which seasonal variation was shared among populations helped inform whether environmental similarity among populations could provide useful predictive information for forecasting epidemic dynamics.

#### Forecasting models

The time series for each forecast variable (disease prevalence, infected host density, and healthy host density) were analysed using three suites of forecasting models: a seasonal naïve benchmark model, an ARIMA model, and a time-series regression model. These three approaches represent increasing levels of model complexity, from a simple seasonal baseline to models that incorporate temporal dependence and environmental predictors.

The benchmark models provided a baseline against which the performance of more complex forecasting approaches could be evaluated. Specifically, I used a seasonal naïve forecasting approach, which assumes that future values will be equal to the observed value from the previous season (Hyndman, 2021). Although this is a simple forecasting method, seasonal naïve models can provide strong predictions when time series exhibit consistent seasonal patterns and therefore represent an appropriate reference point for evaluating more complex models.

The second group of models consisted of ARIMA models, which forecast future values based on temporal patterns within the observed time series, including autoregressive components (relationships with previous observations), differencing to account for non-stationarity, and moving average components (relationships with previous forecast errors). ARIMA models can also incorporate seasonal structure; however, throughout this manuscript I refer to these models as ARIMA rather than SARIMA because model selection was performed using the auto.arima() function in the R package **forecast**, which can select models with or without seasonal components depending on the support provided by the data. Therefore, not all fitted models necessarily included seasonal terms. The auto.arima() function selected the model structure that minimised corrected Akaike information criterion (AICc; Hyndman, 2021), resulting in different combinations of autoregressive, differencing, and moving average terms among populations and forecast variables.

The third group of models used time-series regression to incorporate temperature as an environmental predictor. These models assume that variation in the forecast variable can be partially explained by variation in temperature (Hyndman, 2021). For regression models based on mean population data and temperature-weighted population data, disease prevalence, log-transformed infected host density, and log-transformed healthy host density were modelled as functions of mean temperature. For single-population regression models, each forecast variable was modelled using temperature from the focal population.

For all regression models, temperature observations from the fourth year of the experiment were used as future predictor values when generating forecasts. This represents a realistic forecasting scenario in which environmental conditions can be measured or estimated ahead of time, whereas future disease dynamics remain unknown. Therefore, the regression models used information about expected environmental conditions to improve predictions of future epidemic dynamics, analogous to forecasting scenarios where temperature forecasts are available but disease observations are not.

#### Training and test data

To evaluate whether spatial information can compensate for limited temporal data, I trained forecasting models using three alternative datasets: data from only the focal population (single population models), average epidemic dynamics across all other populations (average population models), and temperature-weighted averages of all other populations, in which populations with temperature dynamics more similar to the focal population contributed more strongly to the forecasts (weighted average population models). Single population models served as the baseline against which the two multi-population approaches were compared.

For all training datasets, model fitting was restricted to the first three years of observations, while forecasts were evaluated using the held-back fourth year. This design mimics a forecasting scenario in which models are trained on historical data and tested on previously unseen observations from a future epidemic season, providing an independent assessment of their predictive performance.

For the mean and temperature-weighted mean training datasets, the time series of disease prevalence, log diseased adult density, and log healthy adult density from all 20 populations were partitioned into every possible combination of one focal (test) population and the remaining 19 training populations (Fig. 1A). This procedure allowed forecast accuracy to be evaluated repeatedly across different focal populations using independent training and test datasets.

**Figure 1.**
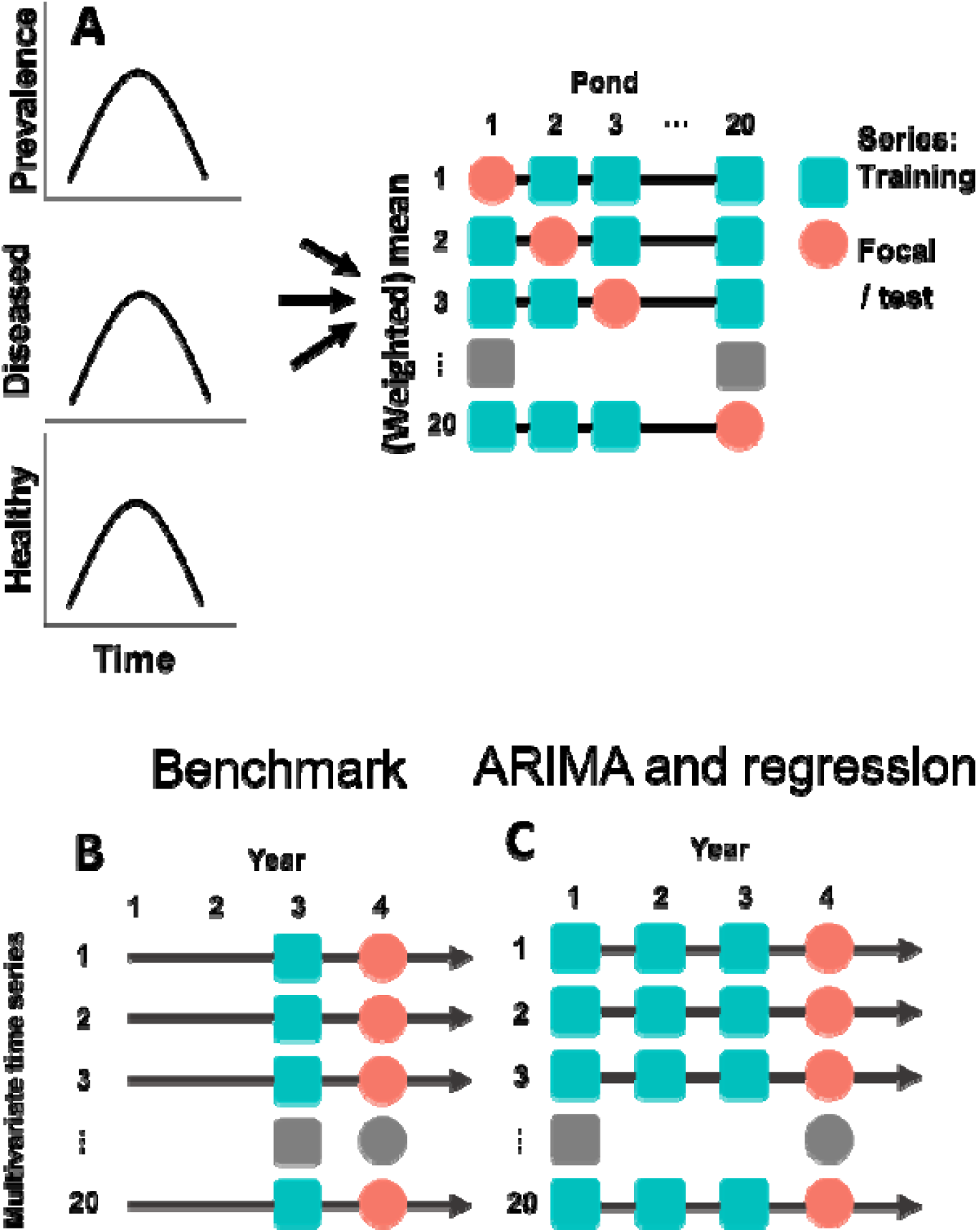
Construction of training datasets and structure of training and test data used for forecasting. Panel A illustrates how the mean and temperature-weighted mean training datasets were calculated, whereas panels B and C illustrate how the different training datasets were used to generate forecasts and evaluate predictions using held-back test data. A) Calculation of mean and temperature-weighted mean training datasets. The time series data for disease prevalence, log diseased adult density, and log healthy adult density from all 20 populations (ponds) were divided into all possible combinations of one focal (test) population and 19 non-focal populations used to construct the training datasets. For each focal population, the mean training dataset was calculated by averaging the response variable across the 19 non-focal populations at each time point. The temperature-weighted mean training dataset was calculated by weighting each non-focal population according to its similarity in seasonal temperature to the focal population. B) Structure of training and test data for benchmark models. For each focal population, forecasts were generated using three training datasets: focal population data, mean population data, and temperature-weighted mean population data. For the mean and temperature-weighted mean datasets, epidemic data from the first three years were combined into a global average time series (blue squares). For focal population datasets, only the third year of epidemic data was used as training data (blue squares), and the fourth year of epidemic data was held back as test data to evaluate forecast accuracy (red circles). C) Structure of training and test data for ARIMA and time-series regression models. For each focal population, the three training datasets consisted of the first three years of epidemic data (blue squares), and the fourth year of epidemic data was used as independent test data (red circles). Unlike the benchmark models, ARIMA and timeseries regression models retained the temporal structure of each training dataset rather than reducing the training data to a global average. For panels B and C, each multivariate time series (MTS) had a frequency of 31 weeks in the first three years and 26 weeks in the final year. The direction of time is represented by black arrows. Grey shapes represent excluded population-years without epidemics.

The mean training dataset was calculated by averaging each response variable across the 19 training populations at each weekly time point. For the temperature-weighted mean dataset, a similarity weight was first calculated for each training population-year as:

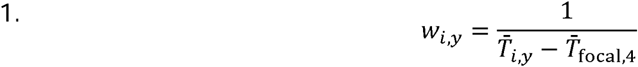

where *w_i,y_* is the temperature similarity weight assigned to training population *i* in year *y*, *T̅_i,y_*, is the seasonal mean temperature of training population *i* in year *y*, and *T̅*_focal,4_, is the seasonal mean temperature of the focal population during the held-out fourth year. Thus, training population-years with seasonal temperature regimes more similar to that of the focal population in the forecast year contributed more strongly to the temperature-weighted mean. A small constant of 0.1 was added to the denominator to avoid division by zero when the seasonal mean temperatures of a training population-year and the focal population were identical.

The temperature-weighted mean training dataset at each weekly time point was then calculated as:

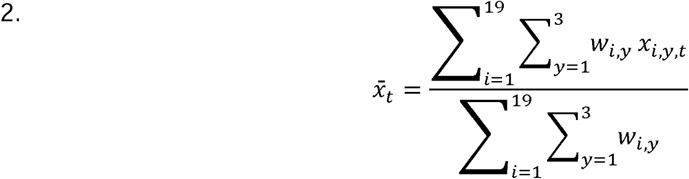

where *x̅_t_* the temperature-weighted mean at week *t*, *x_i,y,t_* is the disease prevalence, log diseased adult density, or log healthy adult density for training population *i,* in year *y,* at week *t*, and *w_i,y_* is the corresponding temperature similarity weight. The small constant had a negligible effect on the resulting weighted mean because it was only applied when seasonal mean temperatures were identical.

Due to the structure of the benchmark model, which was based on a seasonal naïve approach, forecasts for the focal population were generated by repeating its most recently observed seasonal pattern, corresponding to the third year of epidemic data (Fig. 1B). Thus, the focal-population training dataset comprised only the third year rather than an average across the three training years, as averaging across years would not constitute a standard seasonal naïve forecast. In contrast, forecasts based on the mean and temperature-weighted mean training datasets were generated using a single pooled seasonal pattern across the non-focal training populations (Fig. 1B). In contrast, ARIMA and time-series regression models can incorporate temporal autocorrelation and population-specific dynamics, and were therefore fitted separately to each pond’s three-year training time series (Fig. 1C). This distinction allowed the benchmark models to test whether shared epidemic patterns among populations could provide useful forecasts, while the ARIMA and regression models tested whether retaining population-specific temporal structure improved predictive performance.

#### Forecast and error calculation

For all three suites of forecasting models, a forecast horizon of 26 weeks was used to match the time series frequency in the final year of data. Although the dataset contained 20 populations, not all populations experienced an epidemic in every year available for forecasting. Population-years without an epidemic were excluded because disease prevalence and infected host density forecasts could not be evaluated under non-epidemic conditions. This resulted in a final set of 17 forecast occasions for each training data type: focal population, mean population, and temperature-weighted mean population data. To ensure comparability among response variables, the same set of forecast occasions was used for disease prevalence, infected host density, and healthy host density. These same forecast occasions were used in Fig. 2, Fig. S1 and Fig. S2 and all subsequent statistical analyses of forecast error.

**Figure 2.**
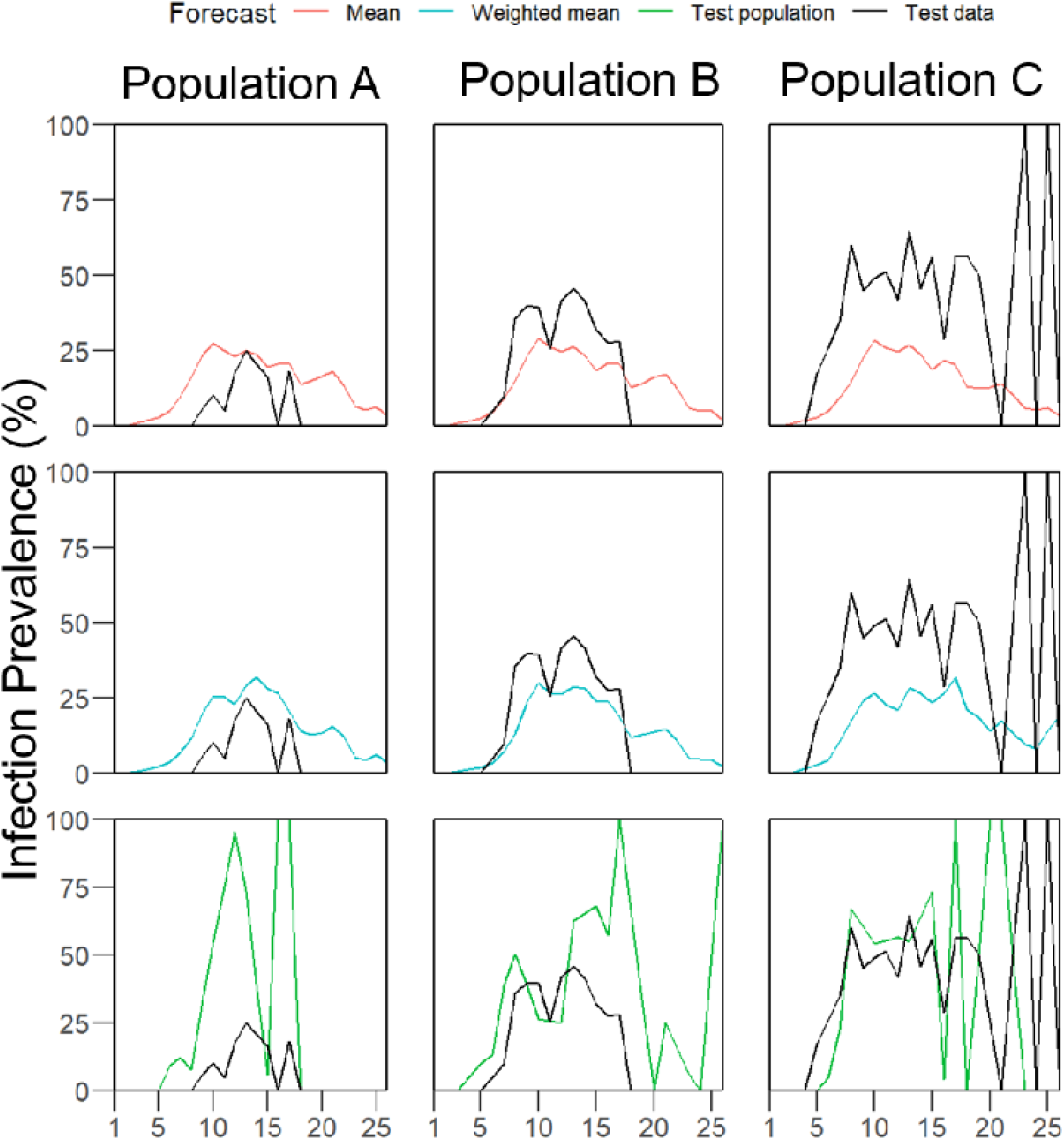
Comparisons of observed and predicted disease prevalence for three representative populations (selected from a total of 20). Observed prevalence (black) is shown alongside forecasts generated by the same benchmark (seasonal naïve) model trained using three different datasets: mean population (red), temperature-weighted mean population (blue), and focal population (green). The complete set of forecast model × training data comparisons is shown in Fig. S1.

Forecast accuracy was determined using root mean squared error (RMSE), which measured the difference between predicted and observed values. For the ARIMA and time series regression models, the forecasts predicted some negative values on a normal scale and some extremely small values on a log scale. These were removed by zero-bounding the disease prevalence and effectively zero-bounding the log diseased and log healthy adult density (log(0.01)-bound). This involved adding the difference between zero and the minimum disease prevalence value to the forecast or adding the difference between log(0.01) and the minimum log diseased or log healthy adult density to the forecast. The average adjusted RMSE was calculated for each set of training data.

To illustrate the forecasting approach, Fig. 2 presents observed and predicted disease prevalence for a subset of three representative populations. The figure compares forecasts generated by the benchmark (seasonal naïve) model using each of the three training datasets (mean population, temperature-weighted mean population, and focal population). The complete set of forecast model and training data comparisons for all forecast variables is provided in Fig. S1. For interpretation of Fig. S1, it is important to note that the auto.arima() procedure selected different model structures for different populations, resulting in variation in ARIMA forecast behaviour. For some populations, the selected model produced forecasts that were nearly identical to the seasonal naïve benchmark, whereas for others the forecasts appeared smoother or more regular because of the fitted autoregressive and moving average structure. These trajectories are statistical forecasts generated from the fitted ARIMA models rather than deterministic simulations.

### Q2: How model and training data affect forecast error

Q2: In general, does forecast error vary by the class of forecast model, the data used to train a model and the temperature of the focal population? This was quantified using separate generalized least squares (GLS) models were fitted to RMSE values for prevalence, infected density, and healthy density, as these represent distinct response variables with different scales and error distributions. Models included forecast model, training data type, and average fourth-year temperature of the focal population as predictors. A heterogeneous variance structure was specified using a variance function that allowed residual variances to differ among training data types.

To assess the overall effects of forecast model, training data type, their interaction, and temperature, nested GLS models with identical variance structures were compared using likelihood ratio tests. These model comparisons were performed using maximum likelihood (ML) estimation, as ML provides valid comparisons among models differing in their fixed-effects structure.

### Q3: Lowest error combinations

Q3: Is forecast error consistently lower for specific combinations of forecast model and training data? This was quantified using final GLS models that were refitted using restricted maximum likelihood (REML) to obtain parameter estimates while accounting for the selected variance structure. To determine whether forecast error was consistently lower for specific combinations of forecast model and training data type, estimated marginal means (EMMs) were calculated from the final GLS models using the emmeans package. Pairwise comparisons were performed among forecast models within each training data type and among training data types within each forecast model. These comparisons were conducted separately for prevalence, infected density, and healthy density RMSE models to identify specific forecast model–training data combinations associated with differences in forecast error while accounting for the fitted GLS model structure and heterogeneous variance among training data types.

## Results

### Seasonal variation

To examine the expected seasonality of the epidemiological variables and assess whether seasonal temperature patterns could underlie these temporal dynamics (Q1), seasonal trajectories of temperature, prevalence, and host densities were plotted together and compared by visual inspection (Fig. 3).

**Figure 3.**
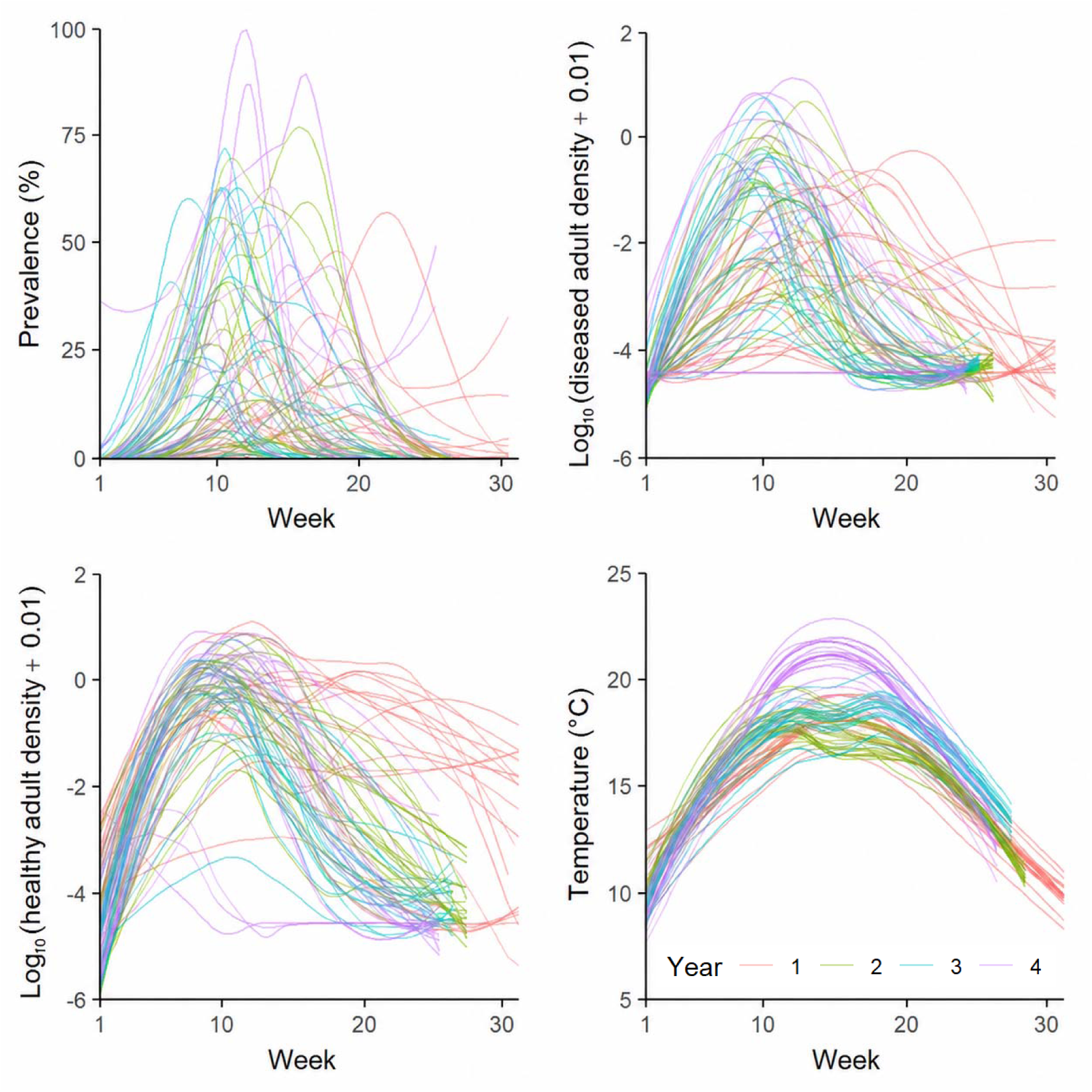
Seasonal trajectories of (A) infection prevalence, (B) log_₁₀_(diseased adult density + 0.01), (C) log_₁₀_(healthy adult density + 0.01), and (D) temperature across all study populations. Lines represent locally weighted regression (LOESS) fits for individual populations. To constrain fitted values to the biologically meaningful range (_≥_ 0%), prevalence was modelled using a log(1 + *x*) transformation prior to LOESS smoothing and back-transformed using the inverse transformation, exp(*x*) − 1. Line colours indicate the year of observation (Years 1–4).

Disease prevalence exhibited pronounced seasonal epidemics across populations, generally increasing rapidly during spring, peaking between weeks 10 and 18, and declining to near zero by the end of the season (Fig. 3). However, both the timing and magnitude of epidemic peaks varied substantially among populations, with maximum prevalence ranging from less than 10% to almost 100%. Diseased adult density followed a similar seasonal trajectory to prevalence, whereas healthy adult density generally increased during early spring before declining as epidemics developed. Although considerable variation existed among populations, these epidemiological variables consistently exhibited strong seasonal dynamics.

Temperature followed a highly consistent seasonal pattern across all populations, increasing from approximately 10°C at the start of the season to peak values of 16–22°C around weeks 12–18 before declining towards the end of the observation period. Notably, peaks in prevalence and diseased adult density broadly coincided with the warmest period of the year, whereas healthy adult density generally declined following peak temperatures. These findings suggest that seasonal temperature may contribute to the timing of epidemic development across populations.

### General predictors of forecast error

To identify general predictors of forecast error (Q2), nested GLS models with identical variance structures were compared using likelihood ratio tests (Table S1). There was no evidence that the effect of forecast model depended on training data type for prevalence (L.R.=5.60, P=0.231), infected density (L.R.=5.66, P=0.226), or healthy density (L.R.=5.60, P=0.231). Forecast model had no significant effect on RMSE for prevalence (L.R.=3.43, P=0.180) or infected density (L.R.=4.29, P=0.117), but significantly affected RMSE for healthy density (L.R.=62.83, P<0.001). Training data type significantly affected RMSE for prevalence (L.R.=11.70, P=0.003), infected density (L.R.=10.36, P=0.006), and healthy density (L.R.=10.58, P=0.005). Temperature significantly improved model fit for prevalence (L.R.=9.60, P=0.002) and infected density (L.R.=17.01, P<0.001), but not healthy density (L.R.=1.15, P=0.283).

### Forecast performance of model and training data combinations

To identify whether specific combinations of forecast model and training data type consistently reduced forecast error (Q3), I used estimated marginal means comparisons of RMSE among forecast models within each training data type and among training data types within each forecast model (Fig. S2; Table S2). Overall, there was limited evidence that a single model–training data combination consistently minimized forecast error across variables.

For prevalence, forecast model performance was generally similar across training data types, with the exception that time-series regression produced higher RMSE than ARIMA when models were trained on focal population data (ARIMA vs. regression: t=2.76, P=0.02). However, comparisons among training data types showed that focal population training resulted in higher RMSE than both mean population and temperature-weighted mean population training for benchmark models (focal vs. mean: t=2.46, P=0.04; focal vs. temperature-weighted mean: t=2.52, P=0.04) and higher RMSE than mean population training for ARIMA models (focal vs. mean: t=2.90, P=0.01). In contrast, regression models showed no differences in RMSE among training data types.

For infected density, model comparisons within training data types revealed no significant differences among forecast models. However, comparisons among training datasets showed that ARIMA models trained on focal population data had higher RMSE than those trained on mean population data (focal vs. mean: t=3.00, P=0.01) and temperature-weighted mean population data (focal vs. temperature-weighted mean: t=3.40, P<0.001). No differences among training data types were detected for benchmark or regression models.

For healthy density, regression models consistently produced lower RMSE than both benchmark and ARIMA models across all multi-population training datasets. Specifically, regression models had lower RMSE than benchmark models when trained on mean population data (benchmark vs. regression: t=−5.23, P<0.001) and temperature-weighted mean population data (benchmark vs. regression: t=−5.26, P<0.001), and lower RMSE than ARIMA models for both mean population (ARIMA vs. regression: t=−5.68, P<0.001) and temperature-weighted mean population training (ARIMA vs. regression: t=−5.64, P<0.001). Regression models trained on focal population data also had lower RMSE than benchmark models (benchmark vs. regression: t=−2.52, P=0.04). Training data type had no effect on RMSE for regression models, but ARIMA models trained using focal population data had higher RMSE than those trained using either mean population data (focal vs. mean: t=3.55, P<0.001) or temperature-weighted mean population data (focal vs. temperature-weighted mean: t=3.51, P<0.001).

Together, these results indicate that the benefits of incorporating data from multiple populations depended on both the forecast variable and model class. Multi-population training improved forecasts primarily for prevalence and for ARIMA forecasts of infected and healthy host densities, whereas regression models generally showed consistent performance across training datasets. The strongest evidence for a consistently superior model–training data combination was observed for healthy host density, where regression models achieved lower RMSE than alternative model classes across multiple training datasets.

## Discussion

Understanding how to forecast infectious disease dynamics is increasingly important for ecological management, yet many natural systems lack the long-term time series required to parameterise reliable forecasting models. I therefore tested whether incorporating information from multiple populations could compensate for limited temporal data and whether weighting populations according to environmental similarity could further improve predictions. Consistent seasonal epidemics were observed across all study populations, with infection prevalence and diseased host density peaking during the warmest part of the year. These seasonal patterns suggested that temperature should provide useful predictive information. However, the forecasting analyses revealed a more nuanced picture. Rather than a single forecasting strategy consistently outperforming all others, forecast accuracy depended on both the response variable and the forecasting approach. Multi-population training improved predictions in some cases, particularly for ARIMA models, whereas regression models consistently produced the most accurate forecasts of healthy host density regardless of the training dataset.

The strong seasonal synchrony between temperature and epidemic dynamics observed across populations suggested that temperature should provide valuable predictive information. Temperature followed a highly consistent seasonal trajectory, with prevalence and diseased host density generally peaking during the warmest weeks of the year, providing biological support for the expectation that models incorporating seasonal temperature—either directly through a temperature predictor or indirectly through seasonal temporal structure—would outperform models that ignored environmental seasonality. However, the forecasting results only partially supported this expectation. Although temperature significantly improved forecasts of prevalence and infected host density, it did not improve forecasts of healthy host density, and incorporating information from multiple populations with similar seasonal timing did not consistently improve prediction accuracy. This suggests that while temperature is an important driver of seasonal epidemics, substantial among-population variation remains that is likely explained by additional ecological factors, such as host demography, parasite transmission dynamics, resource availability, or stochastic processes.

Contrary to my second hypothesis, the results do not support the broad conclusion that multi-population models consistently outperform models trained solely on focal-population data. Nevertheless, they do demonstrate that borrowing information from other populations can improve forecasts under specific circumstances. The clearest example was provided by ARIMA models, for which multi-population training consistently reduced forecast error relative to focal-population training for infected host density and healthy host density, and also improved prevalence forecasts compared with focal-population training in some comparisons. This suggests that when relatively little historical information is available for an individual population, averaging information across replicate populations can provide more stable estimates of the underlying temporal dynamics. By increasing the amount of training data available, multi-population datasets may reduce the influence of stochastic fluctuations or atypical epidemic trajectories present within individual populations.

Interestingly, weighting populations according to similarity in seasonal temperature dynamics did not provide additional improvements beyond simple averaging across populations. This suggests that populations experiencing similar seasonal temperatures did not necessarily exhibit more similar epidemic trajectories than populations selected at random. One explanation is that temperature alone is insufficient to capture the environmental conditions determining epidemic dynamics. Local ecological processes, including variation in host abundance, nutrient availability, parasite introduction timing, or other abiotic conditions, may differ substantially among ponds despite similar thermal regimes (Ebert, 2005; Hall et al., 2011; Lafferty & Holt, 2003). Consequently, weighting populations using temperature similarity alone may fail to identify those with genuinely similar disease dynamics. Future forecasting approaches may therefore benefit from incorporating multiple measures of ecological similarity rather than relying solely on seasonal temperature patterns.

The strongest evidence for differences among forecasting approaches emerged for healthy host density. Regression models consistently outperformed both benchmark and ARIMA models across nearly all training datasets, whereas benchmark and ARIMA models performed similarly for prevalence and infected host density. Unlike prevalence, which is bounded between zero and one, or infected density, which closely tracks epidemic progression, healthy host density reflects the combined effects of host population growth, infection, mortality and demographic turnover (Ebert, 2005). These multiple interacting processes may be better captured by regression models incorporating environmental predictors than by purely autoregressive approaches (Ives et al., 2003). The consistent superiority of regression models therefore suggests that explicitly modelling environmental drivers becomes increasingly important when forecasting demographic variables influenced by multiple ecological processes.

The benefits of borrowing information across populations are likely to depend on ecological similarity among populations. The replicated *Daphnia* system used here represents a relatively controlled case, where shared host–parasite interactions and environmental conditions likely facilitate information transfer. Similar approaches may be most valuable in systems with replicated populations and comparable ecological drivers, such as metapopulations or agricultural disease systems, where long-term data are limited (Liebhold et al., 2006). However, transferability is likely to decline in more heterogeneous systems where differences in habitat, host demography, community interactions, or evolutionary history produce divergent disease dynamics (Altizer et al., 2006). Future forecasting approaches should therefore identify informative populations using multiple ecological traits rather than environmental similarity alone.

Spatial replication may provide a valuable strategy for forecasting emerging diseases where long-term temporal data are unavailable. By generating multiple concurrent epidemic trajectories, replicated systems can help identify general drivers of disease dynamics and reduce reliance on long historical records (Keeling & Rohani, 2008). However, this approach will only improve forecasts when replicated populations share relevant ecological drivers, highlighting the need to combine spatial replication with appropriate measures of ecological similarity. Thus, replicated experiments can complement surveillance efforts by accelerating forecasting during early stages of disease emergence when historical data are limited.

Several limitations should be considered. This study used relatively simple forecasting approaches and temperature as the primary environmental predictor, whereas disease dynamics are often shaped by nonlinear environmental effects, time lags, and interactions among multiple drivers (Cressie & Wikle, 2011; Rohr et al., 2011). Incorporating additional ecological predictors, such as resource availability, host demography, and community context, may therefore improve forecasts by capturing processes that cannot be explained by temperature alone (Hall et al., 2011; Ebert, 2005). Finally, because this study focused on a single host–parasite system in replicated semi-natural ponds, future work should test whether these findings generalise across systems with different transmission modes, ecological complexity, and environmental drivers (Altizer et al., 2006).

Overall, this study demonstrates that replicated populations can provide valuable information for ecological forecasting, but that the benefits of borrowing information across populations are conditional rather than universal. Although the strong seasonal synchrony between temperature and epidemic development suggested that environmentally informed forecasting approaches would consistently outperform simpler alternatives, the empirical results were more complex. Multi-population training improved forecasts primarily for ARIMA models and selected epidemiological variables, whereas regression models provided the clearest and most consistent improvement for healthy host density. These findings highlight the importance of matching forecasting approaches to both the ecological response variable and the available training data, and suggest that exploiting replicated ecological systems offers a promising avenue for improving forecasts when long-term population-specific datasets are unavailable.

## Author contributions

Sam Paplauskas: conceptualization (lead), formal analysis (lead), investigation (lead), methodology (lead), project administration (lead), visualization (lead), writing – original draft (lead), writing – review and editing (lead).

## Acknowledgements

I would like to express my sincere gratitude for the supporting roles played by Drs Stuart Auld and Stephen Thackeray for reviewing early manuscript drafts. Also, big thanks to June Brand for their assistance in collecting data.

## Data availability statement

The data supporting this study are available from the Zenodo repository at https://doi.org/10.5281/zenodo.21411305

## Supporting information

See the supporting information file.

