## Supplemental Figures and Tables for "Borrowing data from other populations to forecast epidemic size"

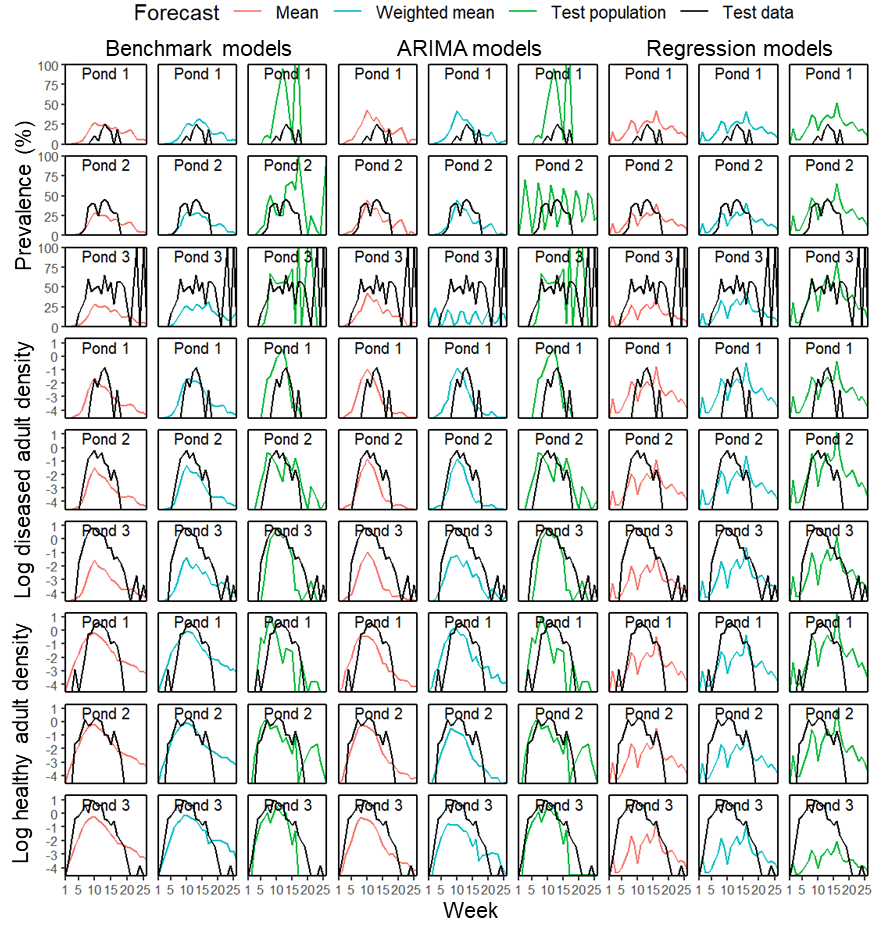

**Figure S1. Variation in forecast accuracy between three example ponds.** For each forecast variable, disease prevalence, log diseased adult density and log healthy adult density, there were three suites of model, benchmark, where the forecast was equal to the values observed in the previous season, auto-regressive integrated moving average (ARIMA) and regression models which used temperature as a predictor. For each model, there were three classes of training data which produced three separate forecasts, the mean, temperature weighted mean and test population forecasts which are indicated by the red, blue and green lines respectively, as well as the corresponding test data which was used to evaluate the accuracy of forecasts as indicated by the black lines. The text shows the different pond numbers.

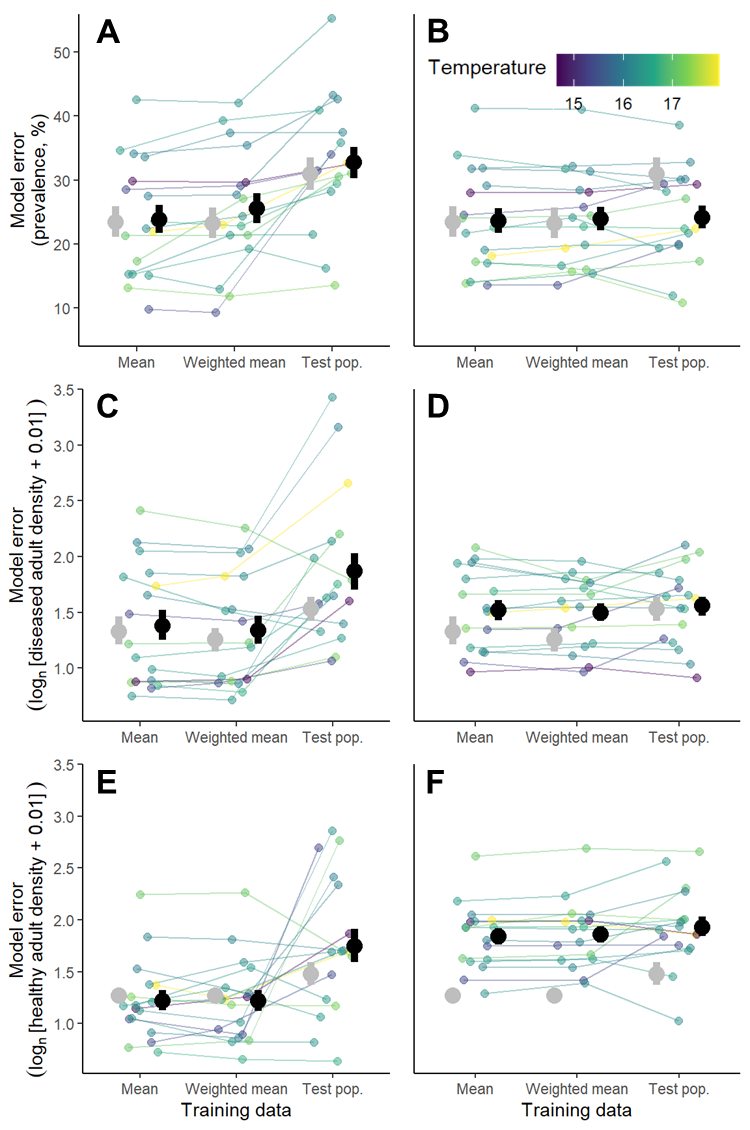

**Figure S2. Model error for forecasts of disease prevalence, log diseased adult density, and log healthy adult density.** Mean model error across the forecast occasions (large points), with error bars indicating standard error, and individual model error for each forecast occasion (small points) are shown. Forecasts were generated from 20 populations; however, population-years without an epidemic were excluded, resulting in 17 forecast occasions used consistently across all three response variables. The colour scale indicates the average temperature of the focal population in year four, and lines connect forecasts from the same population of interest. Results are shown for three sets of models: benchmark (grey points), ARIMA (black points in the first column of panels; A, C, E), and regression (black points in the second column of panels; B, D, F), and three sets of training data: mean population (Mean), temperature-weighted mean population (Weighted mean), and focal population (Test pop.) data. Model error is measured as root mean squared error (RMSE).

Supplementary Table 1. To identify general predictors of forecast error, nested GLS models with identical variance structures were compared using likelihood ratio tests.

| **Response** | **Effect tested** | **L.Ratio** | **P-value** |
| --- | --- | --- | --- |
| Prevalence | Model × Data.type | 5.60 | 0.231 |
| Prevalence | Model | 3.43 | 0.180 |
| Prevalence | Data.type | 11.70 | 0.0029 |
| Prevalence | Temperature | 9.60 | 0.0019 |
| Infected density | Model × Data.type | 5.66 | 0.226 |
| Infected density | Model | 4.29 | 0.117 |
| Infected density | Data.type | 10.36 | 0.0056 |
| Infected density | Temperature | 17.01 | < 0.001 |
| Healthy density | Model × Data.type | 5.60 | 0.231 |
| Healthy density | Model | 62.83 | < 0.001 |
| Healthy density | Data.type | 10.58 | 0.0051 |
| Healthy density | Temperature | 1.15 | 0.283 |

**Table S2. Estimated marginal means comparisons of forecast error (RMSE) among forecast model and training data combinations.**
Pairwise comparisons were conducted using estimated marginal means (*emmeans*). Negative estimates indicate lower RMSE for the first level of the contrast relative to the second level.

**(a) Pairwise comparisons among forecast models within each training data type**

| **Forecast variable** | **Training data type** | **Contrast** | **Estimate** | **SE** | **df** | **t-ratio** | **P-value** |
| --- | --- | --- | --- | --- | --- | --- | --- |
| Prevalence | Test population | Benchmark − ARIMA | -1.78 | 3.09 | 46.42 | -0.58 | 0.83 |
|  |  | Benchmark − Regression | 6.76 | 3.09 | 46.42 | 2.19 | 0.08 |
|  |  | ARIMA − Regression | 8.54 | 3.09 | 46.42 | 2.76 | 0.02 |
|  | Mean population | Benchmark − ARIMA | -0.41 | 3.03 | 42.58 | -0.14 | 0.99 |
|  |  | Benchmark − Regression | -0.19 | 3.03 | 42.58 | -0.06 | 1.00 |
|  |  | ARIMA − Regression | 0.23 | 3.03 | 42.58 | 0.08 | 1.00 |
|  | Temperature-weighted mean | Benchmark − ARIMA | -2.28 | 3.03 | 45.04 | -0.75 | 0.73 |
|  |  | Benchmark − Regression | -0.65 | 3.03 | 45.04 | -0.22 | 0.97 |
|  |  | ARIMA − Regression | 1.63 | 3.03 | 45.04 | 0.54 | 0.85 |
| Infected density | Test population | Benchmark − ARIMA | -0.34 | 0.17 | 47.65 | -1.99 | 0.13 |
|  |  | Benchmark − Regression | -0.03 | 0.17 | 47.65 | -0.16 | 0.99 |
|  |  | ARIMA − Regression | 0.31 | 0.17 | 47.65 | 1.83 | 0.17 |
|  | Mean population | Benchmark − ARIMA | -0.05 | 0.15 | 48.08 | -0.32 | 0.94 |
|  |  | Benchmark − Regression | -0.19 | 0.15 | 48.08 | -1.22 | 0.45 |
|  |  | ARIMA − Regression | -0.14 | 0.15 | 48.08 | -0.90 | 0.65 |
|  | Temperature-weighted mean | Benchmark − ARIMA | -0.09 | 0.14 | 47.42 | -0.62 | 0.81 |
|  |  | Benchmark − Regression | -0.24 | 0.14 | 47.42 | -1.76 | 0.19 |
|  |  | ARIMA − Regression | -0.16 | 0.14 | 47.42 | -1.14 | 0.49 |
| Healthy density | Test population | Benchmark − ARIMA | -0.27 | 0.18 | 47.82 | -1.51 | 0.30 |
|  |  | Benchmark − Regression | -0.45 | 0.18 | 47.82 | -2.52 | 0.04 |
|  |  | ARIMA − Regression | -0.18 | 0.18 | 47.82 | -1.01 | 0.57 |
|  | Mean population | Benchmark − ARIMA | 0.05 | 0.11 | 46.92 | 0.46 | 0.89 |
|  |  | Benchmark − Regression | -0.57 | 0.11 | 46.92 | -5.23 | <0.001 |
|  |  | ARIMA − Regression | -0.62 | 0.11 | 46.92 | -5.68 | <0.001 |
|  | Temperature-weighted mean | Benchmark − ARIMA | 0.04 | 0.11 | 46.81 | 0.38 | 0.92 |
|  |  | Benchmark − Regression | -0.60 | 0.11 | 46.81 | -5.26 | <0.001 |
|  |  | ARIMA − Regression | -0.64 | 0.11 | 46.81 | -5.64 | <0.001 |

**(b) Pairwise comparisons among training data types within each forecast model**

| **Forecast variable** | **Forecast model** | **Contrast** | **Estimate** | **SE** | **df** | **t-ratio** | **P-value** |
| --- | --- | --- | --- | --- | --- | --- | --- |
| Prevalence | Benchmark | Test population − Mean population | 7.52 | 3.06 | 92.77 | 2.46 | 0.04 |
|  |  | Test population − Temperature-weighted mean | 7.71 | 3.06 | 89.99 | 2.52 | 0.04 |
|  |  | Mean population − Temperature-weighted mean | 0.19 | 3.03 | 94.06 | 0.06 | 1.00 |
|  | ARIMA | Test population − Mean population | 8.88 | 3.06 | 92.77 | 2.90 | 0.01 |
|  |  | Test population − Temperature-weighted mean | 7.21 | 3.06 | 89.99 | 2.35 | 0.05 |
|  |  | Mean population − Temperature-weighted mean | -1.67 | 3.03 | 94.06 | -0.55 | 0.85 |
|  | Regression | Test population − Mean population | 0.57 | 3.06 | 92.77 | 0.19 | 0.98 |
|  |  | Test population − Temperature-weighted mean | 0.30 | 3.06 | 89.99 | 0.10 | 0.99 |
|  |  | Mean population − Temperature-weighted mean | -0.28 | 3.03 | 94.06 | -0.09 | 1.00 |
| Infected density | Benchmark | Test population − Mean population | 0.20 | 0.16 | 94.00 | 1.21 | 0.45 |
|  |  | Test population − Temperature-weighted mean | 0.27 | 0.15 | 91.67 | 1.76 | 0.19 |
|  |  | Mean population − Temperature-weighted mean | 0.08 | 0.15 | 94.31 | 0.53 | 0.86 |
|  | ARIMA | Test population − Mean population | 0.48 | 0.16 | 94.00 | 3.00 | 0.01 |
|  |  | Test population − Temperature-weighted mean | 0.53 | 0.15 | 91.67 | 3.40 | <0.001 |
|  |  | Mean population − Temperature-weighted mean | 0.04 | 0.15 | 94.31 | 0.29 | 0.96 |
|  | Regression | Test population − Mean population | 0.04 | 0.16 | 94.00 | 0.23 | 0.97 |
|  |  | Test population − Temperature-weighted mean | 0.06 | 0.15 | 91.67 | 0.38 | 0.92 |
|  |  | Mean population − Temperature-weighted mean | 0.02 | 0.15 | 94.31 | 0.14 | 0.99 |
| Healthy density | Benchmark | Test population − Mean population | 0.21 | 0.15 | 78.14 | 1.39 | 0.35 |
|  |  | Test population − Temperature-weighted mean | 0.21 | 0.15 | 80.26 | 1.42 | 0.34 |
|  |  | Mean population − Temperature-weighted mean | 0.01 | 0.11 | 94.95 | 0.05 | 1.00 |
|  | ARIMA | Test population − Mean population | 0.53 | 0.15 | 78.14 | 3.55 | <0.001 |
|  |  | Test population − Temperature-weighted mean | 0.53 | 0.15 | 80.26 | 3.51 | <0.001 |
|  |  | Mean population − Temperature-weighted mean | 0.00 | 0.11 | 94.95 | -0.01 | 1.00 |
|  | Regression | Test population − Mean population | 0.09 | 0.15 | 78.14 | 0.63 | 0.80 |
|  |  | Test population − Temperature-weighted mean | 0.07 | 0.15 | 80.26 | 0.48 | 0.88 |
|  |  | Mean population − Temperature-weighted mean | -0.02 | 0.11 | 94.95 | -0.20 | 0.98 |

**Note:** RMSE = root mean square error. P-values are adjusted pairwise comparisons from estimated marginal means. “Regression” refers to time-series regression models. “Temperature-weighted mean” refers to population data weighted according to temperature similarity with the focal population.
